# Chimeric anti-HLA antibody receptor engineered human regulatory T cells suppress alloantigen-specific B cells from pre-sensitized transplant recipients

**DOI:** 10.1101/2025.03.27.645777

**Authors:** Jaime Valentín-Quiroga, Alejandro Zarauza-Santoveña, Eduardo López-Collazo, Leonardo M.R. Ferreira

## Abstract

Organ transplantation is a lifesaving procedure, with 50,000 transplants happening every year in the United States. However, many patients harbor antibodies and B cells directed against allogeneic human leukocyte antigen (HLA) molecules, notably HLA-A2, greatly decreasing their likelihood of receiving a compatible organ. Moreover, antibody-mediated rejection is a significant contributor to chronic transplant rejection. Current strategies to desensitize patients non-specifically target circulating antibodies and B cells, resulting in poor efficacy and complications. Regulatory T cells (Tregs) are immune cells dedicated to suppressing specific immune responses by interacting with both innate and adaptive immune cells. Here, we genetically modified human Tregs with a chimeric anti-HLA antibody receptor (CHAR) consisting of an extracellular HLA-A2 protein fused to a CD28-CD3zeta intracellular signaling domain, driving Treg activation upon recognition of anti-HLA-A2 antibodies on the surface of alloreactive B cells. We find that HLA-A2 CHAR Tregs get activated specifically by anti-HLA-A2 antibody-producing cells. Of note, HLA-A2 CHAR activation does not negatively affect Treg stability, as measured by expression of the Treg lineage transcription factors FOXP3 and HELIOS. Interestingly, HLA-A2 CHAR Tregs are not cytotoxic towards anti-HLA-A2 antibody-producing cells, unlike HLA-A2 CHAR modified conventional CD4^+^ T cells. Importantly, HLA-A2 CHAR Tregs recognize and significantly suppress high affinity IgG antibody production by B cells from HLA-A2 sensitized patients. Altogether, our results provide proof-of-concept of a new strategy to specifically inhibit alloreactive B cells to desensitize transplant recipients.

## Introduction

Organ transplantation represents a pivotal advancement in modern medicine, offering a lifeline to thousands of patients suffering from end-stage organ failure. Renal transplant is the most common organ transplant worldwide, according to the latest Global Observatory on Donation and Transplantation (GDOT) report [1; 2].

However, several hurdles remain in the field of organ transplantation. A significant barrier to successful transplantation is immune rejection of the donor organ by the recipient’s immune system [3]. The current standard of care in organ transplant patients involves lifetime multimodal immunosuppressive drug therapy. Broad suppression of the immune suppression by these drugs results in multiple toxicities, including viral infections, nephrotoxicity, neurotoxicity, hyperglycemia, and cancer development [4; 5; 6; 7; 8]. These side effects are especially pernicious in pediatric transplant recipients, who can suffer growth delays, cognitive impairments, and compounded cancer risk due to continued exposure to steroids and neurotoxic drugs during a critical developmental period [9; 10; 11; 12].

Strikingly, 20% of first-time organ recipients and up to 75% of second-time recipients harbor antibodies and B cells directed against allogeneic human leukocyte antigen (HLA) molecules [13; 14; 15]. HLA-A2 is a very common HLA allele group, with reports of 25% of renal transplant recipients in Europe and the United States receiving an HLA-A2 mismatched renal transplant [16; 17]. These HLA sensitized patients face an uphill battle in securing compatible grafts as the risk of antibody-mediated rejection escalates, narrowing the pool of eligible donor organs [18], creating the need for higher doses of immunosuppressants [19], and contributing to chronic rejection [20].

Current desensitization protocols to mitigate the effects of these antibodies lack specificity, targeting total circulating antibodies or B cells, resulting in poor efficacy and unintended complications [19]. It is thus imperative to develop treatments that specifically target the recipient’s alloreactive B cells. Using a cellular approach instead of a broad pharmacological approach could increase efficacy and help prevent non-specific side effects.

Regulatory T cells (Tregs), a subset of CD4^+^ T cells integral to the maintenance of immune tolerance, play a pivotal role in modulating immune responses against self and non-self antigens. These specialized immune cells exert their suppressive functions through various mechanisms, including the inhibition of effector T cell activation and the modulation of B cell responses [21; 22; 23]. Due to these tolerogenic properties, several ongoing trials focus on using Tregs as cellular therapeutics to replace or diminish the dose of immunosuppressive drugs needed to prevent organ transplant rejection [22; 24; 25; 26].

One strategy to reverse HLA pre-sensitization is thus to engineer Tregs to recognize anti- allogeneic HLA B cells and suppress their function, leading to immune tolerance specifically to the target allogeneic HLA molecule without affecting immunity to other antigens. Such an approach can help more patients become eligible for organ transplants and reduce the need for broad immunosuppressive regimens in transplant recipients. To accomplish this, we genetically modified human Tregs with a chimeric anti-HLA antibody receptor (CHAR) consisting of an extracellular HLA-A2 protein fused to a CD28-CD3zeta intracellular signaling domain [27; 28], driving Treg activation upon recognition of anti-HLA-A2 antibodies on the surface of alloreactive B cells, and assessed HLA-A2 CHAR Treg phenotype and suppressive function towards HLA-A2 sensitized patient B cells.

## Materials and Methods

### Molecular Biology

The HLA-A2-CHAR-2A-NGRFt lentiviral plasmid was synthesized by VectorBuilder (Chicago, IL). The construct contained an MND promoter driving the expression of HLA-A2 fused to a CD8 hinge (H), CD28 transmembrane domain (TM), and a CD28–CD3zeta tandem intracellular signaling domain (Figure 1A), followed by a T2A sequence and a truncated nerve growth factor receptor (NGFRt) as a reporter gene, similar to constructs reported in [27; 28]. Lentivirus particles were produced by VectorBuilder and shipped to the laboratory, where they were stored in aliquots at - 80°C until use.

**Figure 1.**
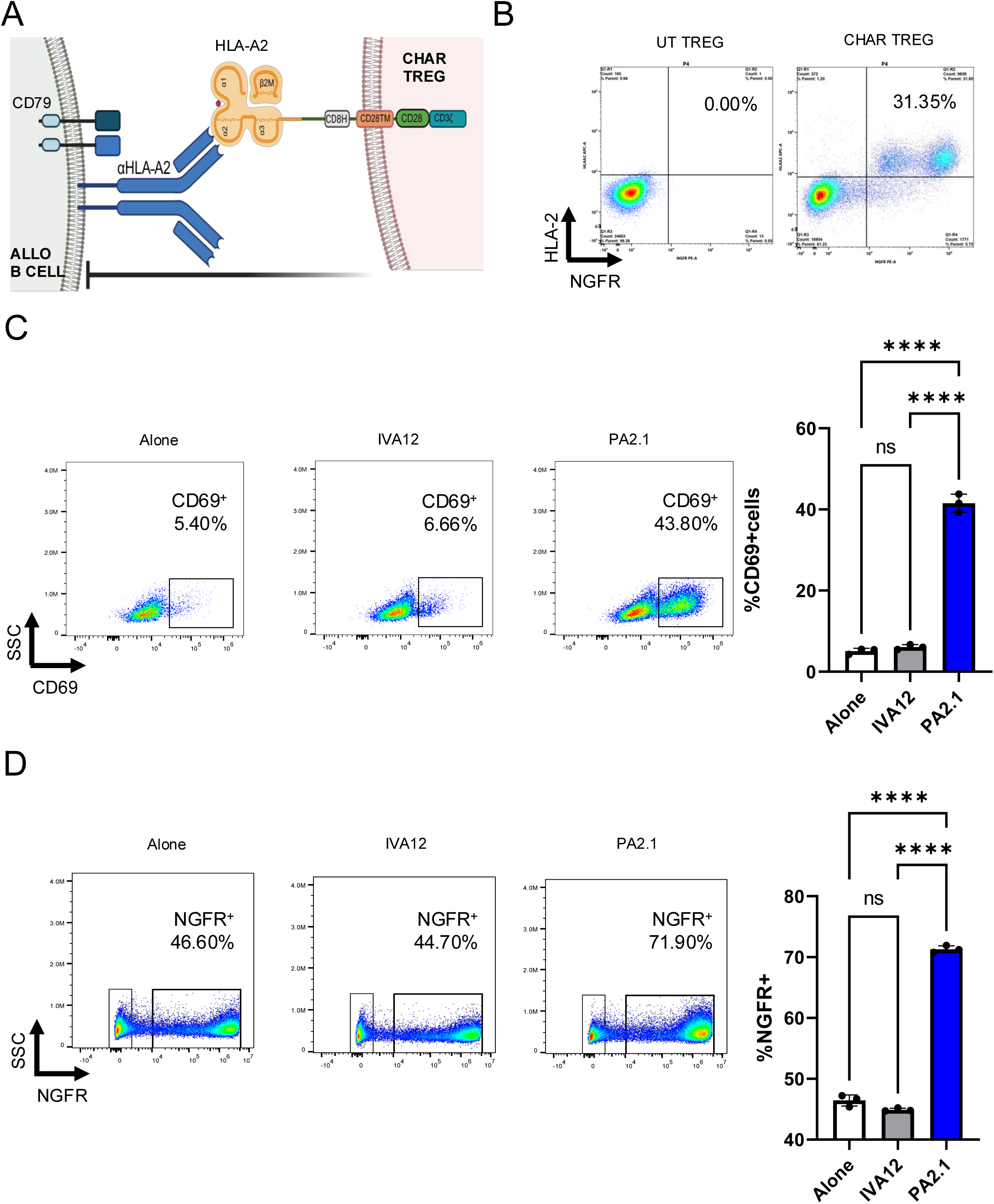
HLA-A2 CHAR Tregs are activated specifically by anti-HLA-A2 antibody producing cells. **(A)** Schematic representation of chimeric anti-human leukocyte antigen (HLA) antibody receptor (CHAR) featuring a CD28-CD3 zeta signaling domain, expressed on the surface of a human regulatory T cell (Treg) binding an anti-HLA-A2 antibody on the surface of an allogeneic B cell from an HLA-A2 pre-sensitized patient. The B cell receptor (BCR) on the surface of a B cell comprises a surface-bound antibody and the signaling heterodimer CD79A and CD79B. Upon engagement, the HLA-A2 CHAR Treg suppresses anti-HLA-A2 expressing B cell function. **(B)** Cell surface expression of HLA-A2 CHAR construct in lentivirus transduced Tregs, as assessed by co-expression of HLA-A2 and a reporter gene, truncated nerve growth factor receptor (NGFRt), linked to the CHAR gene by a 2A peptide. UT, untransduced. **(C)** HLA-A2 CHAR Treg activation, as assessed by surface expression of CD69, upon 48-hour co-incubation with irradiated PA2.1 (anti-HLA-A2) and IVA12 (anti-HLA-DR, DP, DQ) hybridoma cells. **(D)** Enrichment of HLA-A CHAR expressing Tregs upon 9-day co-incubation with irradiated PA2.1 and IVA12 hybridoma cells, as assessed by surface expression of NGFRt reporter. For **(C)** and **(D)**, bars represent mean and standard deviation (n = 3 technical replicates). Data were analyzed by one-way ANOVA with multiple comparisons. ns, not significant; *p < 0.05, **p < 0.01, ***p < 0.001, and ****p < 0.0001.

### Treg sorting, transduction, and expansion

Human Treg isolation, lentiviral transduction, and ex vivo expansion was carried out as previously described [29]. Human peripheral blood leukopaks from de-identified HLA-A2 negative healthy donors were purchased from STEMCELL Technologies (Vancouver, Canada). CD4^+^ T cells were enriched using the EasySep Human CD4+ T Cell Isolation Kit (STEMCELL Technologies), following manufacturer’s instructions. Enriched CD4^+^ T cells were then stained for CD4, CD25, and CD127, and CD4^+^CD25^hi^CD127^low^ regulatory T cells (Tregs), previously shown to be *bona fide* Tregs [30; 31], and CD4^+^CD25^low^CD127^hi^ effector T (Teff) cells were purified by fluorescence-assisted cell sorting (FACS) using a BD FACS Aria II Cell Sorter (Beckton Dickinson, Franklin Lakes, NJ). Post-sort analyses confirmed greater than 99% purity. Tregs were activated in complete medium (RPMI10), comprising RPMI 1640 medium supplemented with 10% fetal bovine serum (FBS), glutamax, penicillin-streptomycin, HEPES, non-essential amino acids (NEAA), and sodium pyruvate (all from Gibco, ThermoFisher Scientific) with anti-CD3/CD28 beads (Gibco, ThermoFisher Scientific) at a 1:1 ratio and 1,000 international units (IU) per ml of recombinant human IL-2 (Peprotech, ThermoFisher Scientific) at 10^6^ cells/ml in 24-wells [32]. 48 hours after activation, Tregs were transduced with CHAR lentivirus at a multiplicity of infection (MOI) of 3 (3 particles per cell) in the presence of IL-2. After adding the lentivirus, Tregs cells were centrifuged at 1,000 g at 32°C for 1 hour. Following transduction, Tregs were maintained and expanded in RPMI10, with fresh medium and IL-2 being given every two days. Tregs received 1,000 IU/ml IL- 2 and CD4^+^ Teff cells received 100 IU/ml IL-2 for 9-12 days. CHAR transduction efficiency was evaluated by flow cytometry based on HLA2 and NGFRt reporter surface expression.

### Activation assay

CHAR Tregs were co-cultured with irradiated (4,000 rad) B-cell hybridoma cell lines (kind gift from Instituto Salud Carlos III) specific for HLA-A2 (PA2.1) or HLA-DR (IVA12) at a 1:1 ratio in RPMI10 medium supplemented with 1,000 IU/ml IL-2 in round-bottom 96-wells. Tregs alone served as a negative control. Surface expression of the T cell activation marker CD69 in CHAR^+^ Tregs was assessed 48 hours later by flow cytometry. Parallel co-cultures were kept for 9 days to assess the frequency of CHAR^+^ Tregs, as measured by surface expression of NGFR, as an additional metric of activation.

### Treg stability assessment

CHAR Tregs were co-cultured with irradiated PA2.1 or IVA12 B-cell hybridoma cell lines at a 1:1 ratio in RPMI10 medium supplemented with 1,000 IU/ml IL-2 in round-bottom 96-wells. Tregs alone served as a negative control. Cells were passaged and expanded and expression of the Treg lineage transcription factors FOXP3 and HELIOS was assessed 9 days later by intracellular staining using the FOXP3/Transcription Factor Staining Buffer Set (eBioscience, ThermoFisher Scientific), according to manufacturer’s instructions. Teff cells were stained for FOXP3 and HELIOS as negative controls.

### Cytotoxicity assay

CHAR Tregs, CHAR Teff cells or their untransduced (UT) counterparts were co-incubated with PA2.1 cells at a 1:1 ratio for 24 hours in round-bottom 96-wells. 50 μl supernatant was then carefully removed and target cell killing assessed using the CyQUANT Cytotoxicity Lactate Dehydrogenase (LDH) Release Assay kit (Thermofisher Scientific) as per manufacturer’s instructions [33].

### Allogeneic B cell stimulation

Peripheral blood mononuclear cells (PBMCs) from de-identified HLA-A2 pre-sensitized patients (Paediatric and Adult Nephrology Unit, La Paz University Hospital, Madrid, Spain) were thawed in RPMI10, counted, and the frequency of B cells (CD19^+^CD20^+^ cells) was determined by spectral flow cytometry. On the same day, PBMCs were incubated with 100 IU/ml IL-2, 100 IU /ml IL-6 (Peprotech, ThermoFisher Scientific) [34], and irradiated (4,000 rad) HLA2-expressing K562 cells (a kind gift from Jack Strominger, Harvard University) at a ratio of 1 B cell : 10 irradiated HLA-A2- K562 cells with or without HLA-A2 CHAR Tregs at a ratio of 1 CHAR Treg : 1 B cell in 12-wells for up to 5 days. PBMCs from pre-sensitized patients with IL-2 and IL-6 alone, as well as PBMCs from a de-identified healthy donor (STEMCELL Technologies) subjected to all 3 conditions (Figure 3B), were kept as negative controls.

### IgG antibody production assay

Three conditions were set up with HLA-A2 pre-sensitized patient PBMCs and healthy donor PBMCs: PBMCs alone (with IL-2 and IL-6), PBMCs with irradiated HLA2-K562 cells, and PBMCs with irradiated HLA2-K562 cells and HLA-A2 CHAR Tregs, as described above (Figure 3B). 250 μl supernatant was collected from each allogeneic B cell stimulation condition at 48h and 5 days and diluted 1:2 with PBS. Human IgG solid-phase sandwich enzyme-linked immunosorbent assay (ELISA) (Thermofisher Scientific) was performed as per manufacturer’s instructions.

### Spectral flow cytometry

Spectral flow cytometry data was acquired in a 3-laser Cytek Northern Lights spectral flow cytometer (Cytek Biosciences, Fremont, CA). Spectroflow 3.2.1 (Cytek Biosciences) and FlowJo v10.9 software (BD Life Sciences, Franklin Lakes, NJ) were used for flow cytometry data analysis. Data were manually pre-gated to remove cell aggregates, dead cells, debris, and then sub- sampled to include 10,000 live singlets from each sample. Uniform Manifold Approximation and Projection (UMAP) analysis was performed to visualize the different sub-populations in groups [35]. UMAP settings were as follows: all files used, all compensated fluorescent parameters were used besides Live/Dead,Neighbors =15, Minimum Distance = 0.5, Components = 2, Metric = Euclidean. Antibodies used for spectral flow cytometry in this study can be found in Supplementary Table 1.

## Results

Aiming to test the concept of regulatory T cell (Tregs) engineered to specifically recognize and inhibit alloreactive B cells responsible for human leukocyte antigen (HLA) pre-sensitization in patients, we constructed a chimeric anti-HLA antibody receptor and UT (CHAR) specific for anti-HLA-A2 B cells by fusing an HLA-A2 molecule to a CD28 hinge, a CD28 transmembrane domain, and a CD28-CD3zeta intracellular signaling domain [27; 28] (Figure 1A). We then sorted CD4^+^CD25^hi^CD127^low^ Tregs [30; 31] from human peripheral blood collected from HLA-A2 negative donors, activated them with anti-CD3/CD28 beads and IL-2, and transduced them two days later with lentivirus coding for the HLA-A2 CHAR. Flow cytometry analysis of transduced Tregs revealed successful expression of the CHAR construct, as assessed by simultaneous surface expression of HLA-A2 and the truncated nerve growth factor receptor (NGFRt) reporter gene, linked to CHAR gene expression via a 2A peptide (Figure 1B).

To evaluate CHAR target recognition, we co-incubated CHAR Tregs with B-cell hybridoma cell lines specific for HLA-A2 (PA2.1) or for HLA-DR (IVA12). CHAR Tregs upregulated surface expression of the T cell activation marker CD69 (Figure 1C) and increased in frequency (Figure 1D) upon co-incubation with PA2.1 cells, but not with IVA12 cells, indicating CHAR Treg reactivity specifically to HLA-A2 antibody-producing cells.

Next, we sought to confirm that activation via the CHAR did not compromise Treg identity stability, measured by the expression level of the Treg lineage transcription factors FOXP3 and HELIOS (REFs). We found no difference in the frequency of FOXP3 or HELIOS expressing cells between untransduced (UT) or CHAR Tregs co-incubated with either PA2.1 or IVA12 hybridoma cells (Figure 2A). Of note, while the frequency of FOXP3^+^ CHAR Tregs (Figure 2B) and FOXP3^+^HELIOS^+^ CHAR Tregs (Figure 2D) did not change across conditions, FOXP3 levels were higher in CHAR Tregs co-incubated with PA2.1 cells (Figure 2C, an additional line of evidence for CHAR Treg activation specifically by anti-HLA-A2 antibody-producing cells.

**Figure 2.**
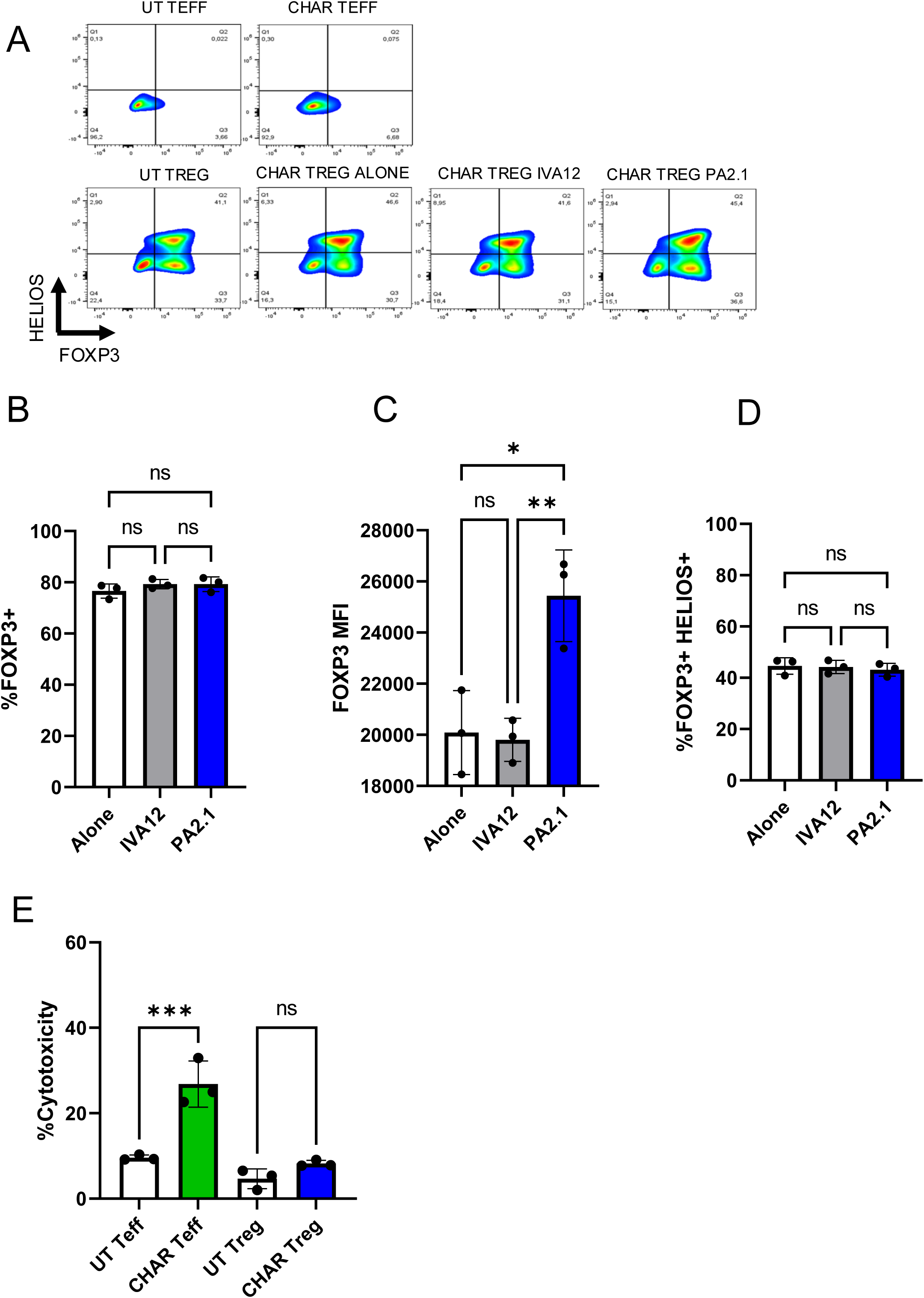
HLA-A2 CHAR Tregs remain stable and are not cytotoxic upon activation. **(A)** Representative flow cytometry analysis of CHAR Treg FOXP3 and HELIOS expression after 9 days of co-culture with irradiated PA2.1 (anti-HLA-A2) and IVA12 (anti-HLA-DR, DP, DQ) hybridoma cells. Untransduced (UT) T effector (Teff) cells and CHAR Teff cells were used as negative controls. **(B)** Frequency of FOXP3^+^ CHAR Tregs alone or co-cultured with PA2.1 and IVA12 hybridoma cells for 9 days. **(C)** FOXP3 expression (mean fluorescence intensity - MFI) in CHAR Tregs alone or co-cultured with PA2.1 and IVA12 hybridoma cells for 9 days. **(D)** Frequency of FOXP3+HELIOS+ CHAR Tregs alone or co-cultured with irradiated PA2.1 and IVA12 hybridoma cells for 9 days. **(E)** Cytotoxicity of HLA-A2 CHAR Tregs and HLA-A2 CHAR Teff cells towards PA2.1, as measured by lactate dehydrogenase (LDH) release after 24-hour co- incubation at a 1:1 ratio. Bars in (B), (C), (D), and (E) represent mean and standard deviation (n = 3 technical replicates). Data were analyzed by one-way ANOVA with multiple comparisons. ns, not significant; *p < 0.05, **p < 0.01, ***p < 0.001, and ****p < 0.0001.

Interestingly, CHAR Tregs were not cytotoxic towards anti-HLA-A2 antibody-producing PA2.1 cells at a 1:1 ratio, unlike CHAR T effector (Teff) cells (Figure 2E), further confirming that HLA-A2 CHAR activation does not destabilize Treg identity.

To assess CHAR Treg function, we thawed peripheral blood mononuclear cells (PBMCs) from HLA-A2 pre-sensitized patients (SEN) and a healthy donor (HD) (Figure 3A) and co-incubated them with irradiated HLA-A2-expressing K562 cells as a source of HLA-A2 antigen in the presence of IL-2 and IL-6 with or without CHAR Tregs for 2 days or 5 days (Figure 3B). In two out of three HLA-A2 pre-sensitized patients tested, CHAR Tregs significantly decreased IgG antibody production by the patient’s cells after 48h of co-incubation (Figure 6), demonstrating the ability of CHAR Tregs to inhibit alloreactive B cells.

**Figure 3.**
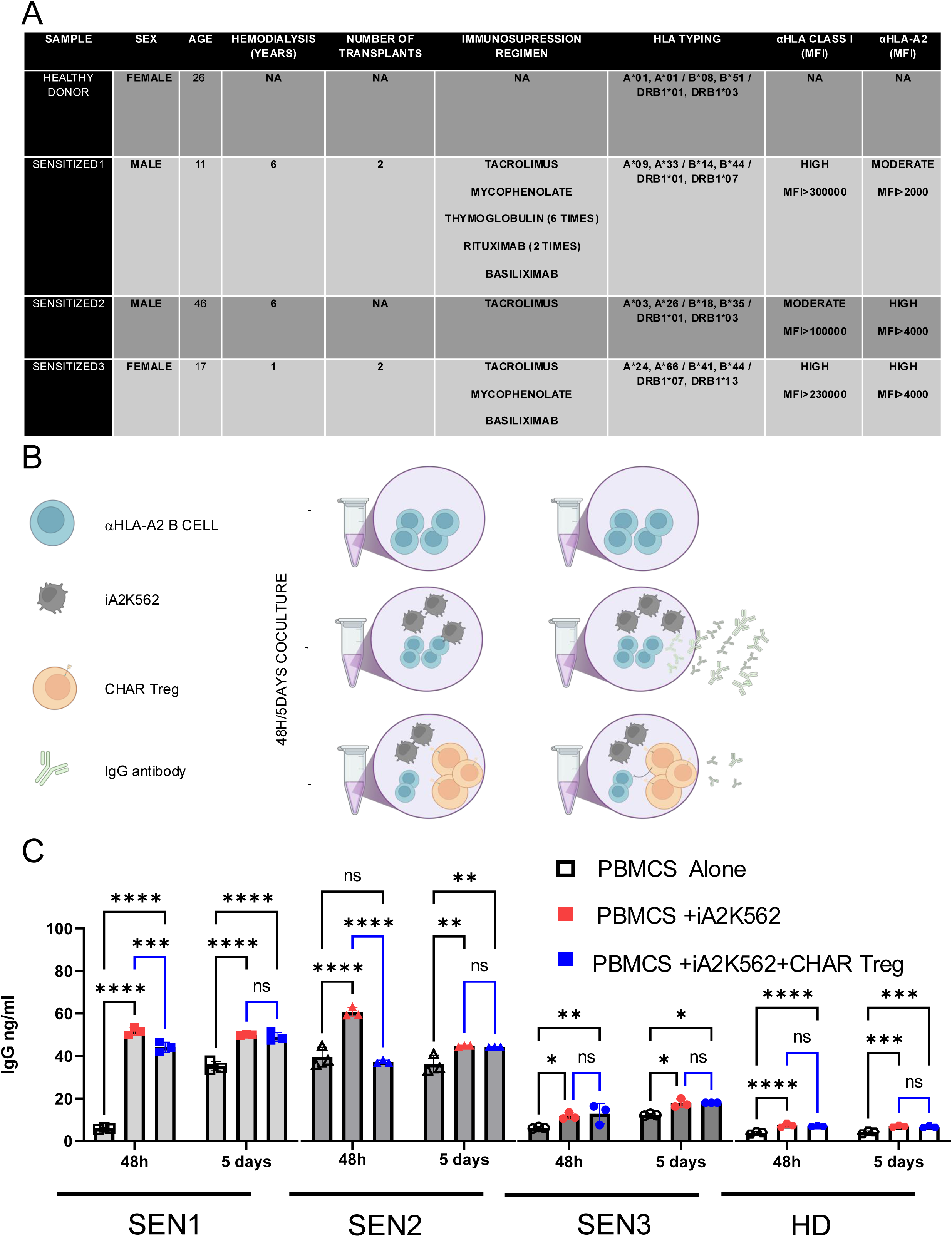
HLA-A2 CHAR Tregs suppress IgG specific production on highly pre sensitized patients. **(A)** Experimental design for assessing HLA-A2 CHAR Treg function in the presence of HLA-A2 pre-sensitized patient cells. HLA-A2 sensitized donor-derived peripheral blood mononuclear cells (PBMCs) were co-incubated with HLA-A2-expressing K562 cells to induce expansion of anti-HLA-A2 B cells and anti-HLA-A2 IgG antibody production. If HLA-A2 CHAR Tregs are added, a decrease in antibody production elicited by exposure to HLA-A2 is expected. **(B)** IgG antibody production 48 hours or 5 days after pre-sensitized patient PBMC co-incubation with HLA-A2-K562 in the presence or absence of HLA-A2 CHAR T regs, as assessed by ELISA (n=3 sensitized patients (SEN) and n=1 healthy donor (HD) control). Data were analyzed by one- way ANOVA with multiple comparisons. ns, not significant; *p < 0.05, **p < 0.01, ***p < 0.001, and ****p < 0.0001. **(C)** Patient and healthy control demographics and clinical characteristics.

In order to gain mechanistic insight into CHAR Treg function, we performed spectral flow cytometric analysis of these co-cultures. We identified NGFR+HLA-A2+CD19- CHAR Tregs and NGFR-HLA-A2-CD19+CD20+ patient B cells, which we then subsetted into naive (CD27^-^IgD^+^), marginal zone (CD27^+^IgD^+^), memory (CD27^+^IgD^-^), and CD27^-^IgD^-^ B cells (Figure 4A). Uniform Manifold Approximation and Projection (UMAP) visualization of total live cells in the different co-culture conditions across all three pre-sensitized patients at 48 hours (Figure 4B) and 5 days (Figure 4D) illustrates that B cells constitute a small and relatively uniform fraction of the patients’ PBMCs (colored in blue, light green, dark green, and orange according to B cell subset) and that CHAR Tregs (colored in red) form distinct clusters, potentially reflecting differences in activation status. Focusing on the B cell fraction, we found that CHAR Tregs significantly reduced the frequency of total B cells in all three pre-sensitized patients’ PBMCs at 48 hours (Figure 4B) and at 5 days (Figure 4D) of co-culture. Of note, all B-cell subsets measured (naïve, memory, marginal zone, and CD27^-^IgD^-^ B cells) were equally depleted by HLA-A2 CHAR Tregs at both time points (Figure 4B, 4D). With regards to the CHAR Tregs, we observed that the majority of CHAR Tregs expressed high levels of CD27, a marker associated with Treg suppressive function [36] after 48h (Figure 4C) and 5 days (Figure 4E) of co-culture. There was a trend where a larger fraction of CHAR Tregs expressed CD27 when co-incubated with SEN PBMCs (86% at 48 hours, 76-88% at 5 days) than with HD PBMCs (80% at 48 hours, 68% at 5 days) (Figure 4C, 4E). Moreover, CHAR Tregs were activated, as assessed by CD71 upregulation [29], in the presence of all three SEN PBMCs, but not HD PBMCs after 48 hours (Figure 4C) and 5 days (Figure 4E) of co-culture.

**Figure 4.**
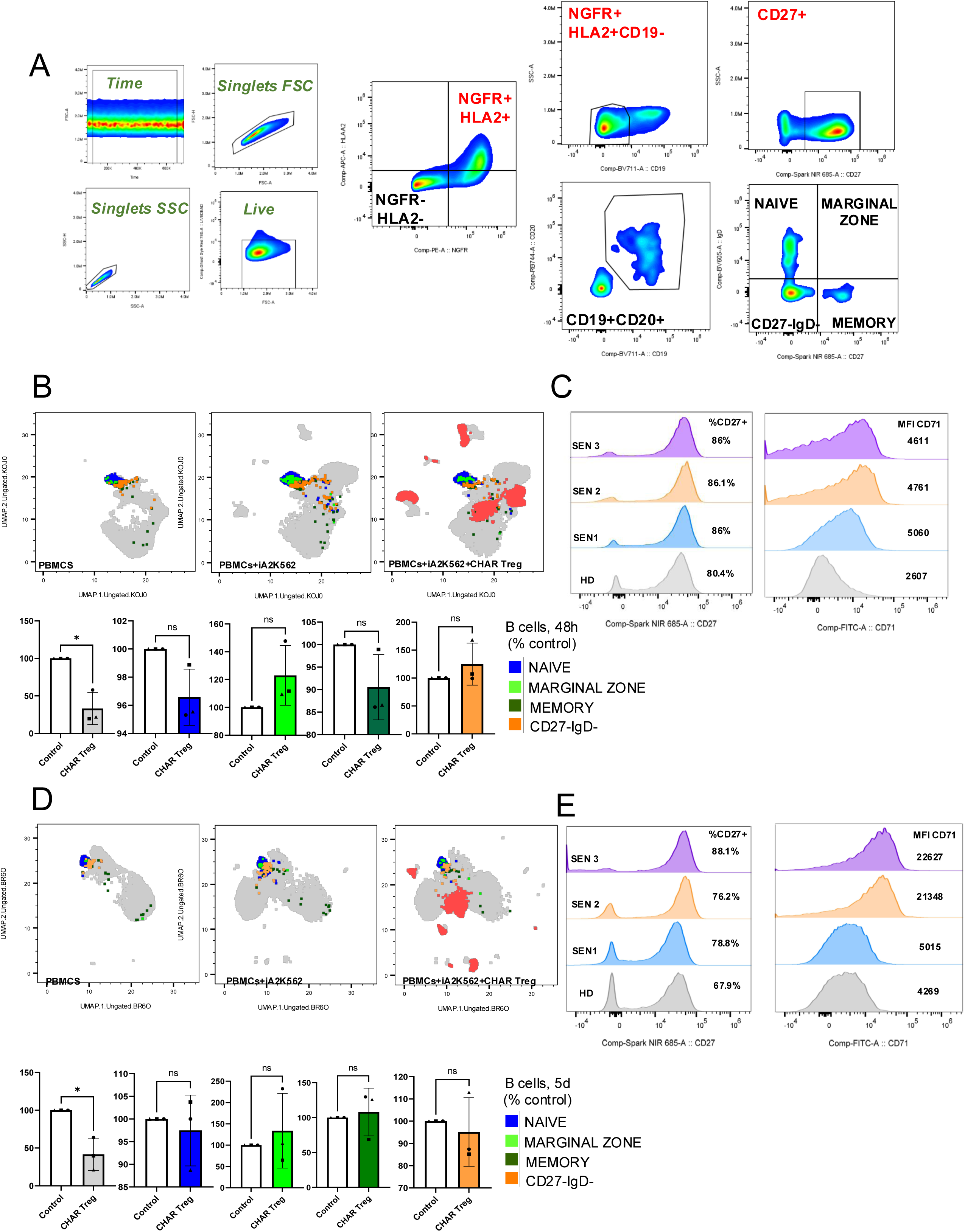
HLA-A2 CHAR Tregs were activated and reduced the frequency of total B cells in pre-sensitized patient peripheral blood samples. **(A)** Flow cytometry gating strategy to identify HLA-A2^+^NGFR^+^CD19^-^ CHAR Tregs, CD19^+^CD20^+^ B cells, and B cell subsets in co-cultures of sensitized patient PBMCs with irradiated HLA-A2-K562 and HLA-A2 CHAR Tregs. **(B,D)** Uniform manifold approximation and projection (UMAP) representation of sensitized patient PBMCs with irradiated HLA-A2-K562 (iA2K562) and HLA-A2 CHAR Tregs depicting naive, marginal zone, memory and IgD^-^CD27^-^ B cells after 48 hours **(B)** and 5 days **(D)** of co-culture. Red subset represents HLA-A2 CHAR Treg in all UMAPs. **(C,E)** Frequency of CD27 expression (left) and CD71 surface expression levels (right) in HLA-A2 CHAR Tregs co-incubated with sensitized patient PBMCs for 48 hours **(C)** and 5 days **(E)**. n=3 sensitized patients (SEN) and n=1 healthy donor (HD) control. Data were analyzed by one-way ANOVA with multiple comparisons. ns, not significant; *p < 0.05, **p < 0.01, ***p < 0.001, and ****p < 0.0001.

## Discussion

The field of immunotherapy has witnessed remarkable advancements in recent years, particularly concerning the engineering of T cells to target specific cells and immune responses. Previous studies have demonstrated the potential of conventional T cells to recognize and eliminate B cells. In the clinic, total B cells are being eliminated using CD19 chimeric antigen receptor (CAR) T cells in patients with systemic lupus erythematosus and other autoimmune disorders, leading to disease remission [37; 38]. At the pre-clinical stage, self-reactive antigen-specific B cells have been eliminated through chimeric autoantibody receptor T cells (CAAR T cells) in the setting of pemphigus vulgaris [39] and alloreactive HLA-specific B cells have been targeted using chimeric HLA receptor (CHAR) T cells [27; 28].

Our study introduces a novel approach by engineering regulatory T cells (Tregs), immune cells dedicated to inhibiting immune responses to maintain immune homeostasis [21; 22; 23], to recognize and suppress B cells specific for an HLA molecule with a CHAR. This innovative strategy not only enhances the specificity of Treg-mediated suppression of alloreactive B cells but also opens new avenues for therapeutic intervention in transplant immunology. Tregs, like all CD4^+^ T cells, possess T cell receptors (TCRs) restricted to human leukocyte antigen (HLA) class II molecules, which are expressed in professional antigen presenting cells, including B cells [40]. Previous work reported the expansion of rare recipient alloreactive Tregs using donor-derived B cells [41]. Yet, antigen-specific Tregs are rare and prone to destabilization upon multiple rounds of activation *ex vivo*, kindling interest in genetic engineering approaches to confer desired antigen specificities to human Tregs [21; 22]. Previous work has focused on modifying human Tregs with artificial receptors to confer them specificity towards transplanted tissues, aiming to provide localized protection from immune attack [42; 43; 44; 45; 46]. Engineering Tregs with CHAR tackles the problem of allogeneic immune rejection from a different, potentially orthogonal and synergistic, angle, providing a targeted mechanism to directly inhibit B cell responses against allogeneic HLA molecules, such as HLA-A2. Unlike engineered conventional T cells, which can induce wide-ranging immune activation and potential tissue damage [47; 48; 49; 50], HLA-A2 CHAR Tregs are designed to suppress only B cells producing anti-HLA-A2 antibodies in an anti-inflammatory fashion, thereby minimizing collateral damage to other components of the immune system or any tissues. In line with this concept, we observed that conventional HLA-A2 CHAR T cells were cytotoxic towards anti-HLA-A2 antibody-producing hybridoma cells, whereas HLA-A2 CHAR Tregs were not (Figure 2E). This specificity and safety profile is crucial in the context of transplantation, where current desensitization strategies often fall short due to their non-specific nature and associated complications [19]. By effectively reducing IgG antibody production in sensitized individuals (Figure 3B), CHAR Tregs offer a promising therapeutic option for improving transplant outcomes by enhancing graft acceptance and reducing the risk of rejection.

Engineering of Tregs to bind specific B cells had been previously demonstrated with FVIII B cell antibody receptor (BAR) Tregs, aimed at inhibiting anti-FVIII antibody production by hemophilic patients treated with recombinant FVIII protein [51]. While this work demonstrated the possibility of directing Tregs towards B cells based on B-cell antigen specificity, our findings extend the applicability of B-cell-targeting engineered Tregs to graft-vs-host disease, organ transplantation, and beyond, potentially addressing challenges in conditions such as miscarriage, where pregnant women develop antibodies against paternally derived HLA molecules [52].

The novelty of our study lies not only in the engineering of Tregs with CHAR but also in the demonstration of their efficacy with HLA sensitized patients’ cells. The ability of CHAR Tregs to suppress IgG production by B cells from pre-sensitized individuals upon exposure to antigen-expressing cells (Figure 3B) is a significant step forward, suggesting that this approach could pave the way for tailored strategies that address individual HLA sensitized patient needs. Polyclonal Tregs have been shown to be safe in phase I and phase II clinical trials [53; 54; 55], and human CAR Tregs targeting transplanted tissues have shown efficacy and safety in preclinical studies [44; 56; 57; 58] and are being tested in ongoing phase I clinical trials [59]. Hence, CHAR Tregs are a good candidate for first-in-human trials, to be used either in pre-sensitized patients with the goal of bringing them to the same baseline as non-sensitized patient and/or as an adjuvant to reduce the doses of immunosuppressive drugs taken by transplant recipients.

Interestingly, CHAR Tregs significantly inhibited IgG production by HLA-A2 antigen-stimulated B cells for only two out of three sensitized patients (Figure 3B). Moreover, CHAR Tregs retained high expression of the activation marker CD71 after 5 days of co-culture also only with two out of three sensitized patient PBMCs (Figure 4E). Future studies are warranted to define what patient characteristics indicate responsiveness to CHAR Treg targeting, as well as to expand the concept to other HLA class I and class II molecules and dissect the mechanisms by which CHAR Tregs inhibit target B cells.

In conclusion, we provide compelling proof-of-concept for a novel immunotherapeutic strategy to desensitize transplant recipients with HLA sensitization. Engineering CHAR Tregs represents an advancement in the quest for specific and safe immunotherapies to modulate harmful B-cell responses in sensitized transplant recipients and beyond. As we continue to refine these technologies, the potential for engineered Tregs to transform clinical practice becomes a tantalizing prospect.

## Data availability statement

The raw data supporting the conclusions of this article will be made available by the authors, without undue reservation.

## Author contributions

JVQ designed the project, designed experiments, conducted experiments, analyzed data, and wrote the manuscript. AZS provided samples and analyzed data. ELC analyzed data. LMRF designed the project, supervised work, designed experiments, conducted experiments, analyzed data, and wrote the manuscript. The manuscript was edited and approved by all authors.

## Acknowledgements

Artwork in certain figures was created using BioRender.com. Thanks are due to Ferreira Lab members for helpful discussions, Instituto Salud Carlos III for kindly gifting the PA2.1 and IVA12 hybridoma cell lines, Jack Strominger (Harvard University) for kindly gifting the HLA-A2-K562 cell line, and Jacob Kendrick, Kirsten Hughes, and Josh Monts at the Flow Cytometry and Cell Sorting Shared Resource, Hollings Cancer Center, Medical University of South Carolina, for excellent technical assistance.

