## Supplemental Table 1 for "Chimeric anti-HLA antibody receptor engineered human regulatory T cells suppress alloantigen-specific B cells from pre-sensitized transplant recipients"

**Supplementary Table 1: Spectral flow cytometry antibodies used in this study.**

| **Antigen** | **Fluorophore** | **Clone** | **Company** |
|---|---|---|---|
| CD4 | Pacific Blue | SK3 | Biolegend,San Diego CA |
| CD19 | BV711 | HIB19 | Biolegend,San Diego CA |
| CD20 | RB744 | 2H7 | BD Biosciences,San Jose,CA |
| CD24 | BV510 | ML5 | Biolegend,San Diego CA |
| CD25 | APC | BC96 | Biolegend,San Diego CA |
| CD27 | Spark NIR 685 | O323 | Biolegend,San Diego CA |
| CD38 | Spark Violet 423 | HIT2 | Biolegend,San Diego CA |
| CD69 | Pe/Cy7 | FN50 | Biolegend,San Diego CA |
| CD71 | FITC | CY1G4 | Biolegend,San Diego CA |
| FOXP3 | PeCy5.5 | PCH101 | eBioscience,San Diego CA |
| HELIOS | FITC | 22F6 | Biolegend,San Diego CA |
| HLA-A2 | APC | BB7.2 | Biolegend,San Diego CA |
| IgD | BV605 | IA6-2 | Biolegend,San Diego CA |
| NGFR | APC | ME20.4 | Biolegend,San Diego CA |
| NGFR | PE | ME20.4 | Biolegend,San Diego CA |
| Ghost Dye | Red 780 | Viability | Tonbo Bioscience,San Diego CA |
